# High-Throughput Screening of FRET-Based Protein Rulers Using a Hyperspectral Microcapillary Array

**DOI:** 10.1101/2025.08.25.672123

**Authors:** Khushank Singhal, Thomas M. Baer, Benjamin D. Allen, Melik C. Demirel

## Abstract

FRET (Fluorescence Resonance Energy Transfer) is a versatile technique used in biology, chemistry, and materials science that involves distance-dependent energy transfer between light-sensitive molecules. Current FRET screening methods are limited when handling large libraries. Challenges include a low dynamic range, overlapping spectra, multiple fluorophores, and high background signals. A high-throughput FRET approach enables rapid, multiplexed screening of interactions, conformations, and dynamics with accurate, real-time measurements. We developed a hyperspectral high-throughput microcapillary array (HyCAP), which is ideal for sorting large clone libraries based on fluorescence assays. Our platform provides sensitive, high-resolution hyperspectral imaging with a throughput of approximately 10^5^ per array. FRET protocols can be used to study variations in protein structure and assemblies. HyCAP demonstrates potential in fluorophore engineering, including emission changes related to environment (such as pH, enzymatic activity, and temperature), stoichiometry differences, fluorophore orientation, and photobleaching through time-resolved experiments.

## INTRODUCTION

Proteins are dynamic macromolecules that perform a vast array of functions in living organisms, including catalyzing metabolic reactions, replicating DNA, responding to stimuli, and transporting molecules, as well as creating structural integrity.^1^ Fluorescent proteins are vital tools in biological screening, widely utilized for applications that range from protein tracking to functional activity assays.^2^ Their key characteristics—excitation and emission spectra, fluorescence intensity, and spectral shape—are essential for accurate and sensitive detection. Over the past decade, extensive efforts in protein engineering have led to the creation of enhanced fluorophores with increased brightness, photostability, and spectral diversity.^3^ Notable examples include mNeonGreen (mNG)^4^, engineered for high brightness and quick maturation, and mScarlet-I (mSc)^5^, developed for its superior red fluorescence and monomeric properties. These engineered fluorophores have significantly broadened the capabilities of fluorescence-based assays, enabling more complex and multiplexed experiments. ^6^

FRET (Fluorescence Resonance Energy Transfer) is a versatile technique used in biology, chemistry, and materials science, which is a distance-dependent energy transfer process between two light-sensitive molecules.^7^ In biology, it is utilized for studying protein interactions (such as receptor-ligand binding and dimerization^8^), protein conformational changes (like enzyme activation and folding^9^), and live-cell imaging (including cellular signaling and protein dynamics^10^). In materials science, it is employed to create protein-based materials (such as self-healing ^11^ and sensors^12^), analyze biopolymer dynamics, and investigate interactions between biomolecules and inorganic nanomaterials.

Despite these advances, current FRET-based screening techniques remain limited in their ability to accurately and efficiently analyze large libraries or sequence variants. Plate readers, while widely used, rely on either fixed filter sets or monochromators, which restrict their spectral resolution and throughput. Flow cytometers, although significantly higher in throughput, typically use multiband filters and cannot capture full spectral profiles, limiting their utility in applications like fluorophore engineering and FRET, where precise spectral information is essential. A high-throughput FRET approach significantly advances protein research by enabling rapid screening of interactions, conformations, and dynamics; multiplexed analysis of multiple protein states; precise quantitative measurements; and real-time observation of protein behavior. Moreover, low dynamic range, difficulty deconvolving overlapping spectra, identifying multiple fluorophores, and high background signals present challenges for FRET. ^13^ Using hyperspectral imaging and high-throughput methods to measure FRET in cell populations or single cells with spectral spectroscopic or microscope systems can address these issues.

Hyperspectral imaging captures many spatial images across numerous closely spaced spectral bands at the same time, with each pixel containing a complete spectrum. It is a highly effective method for characterizing and analyzing biological samples. ^14^ Hyperspectral imaging captures spectral data at each pixel by dispersing light. Specifically, visible spectroscopy uses a grating to separate different frequencies of radiation leaving the sample. This method helps deconvolve overlapping spectra, identify multiple fluorophores, and enhance FRET quantification.

Here, we developed a hyperspectral microcapillary imaging platform called HyCAP to create and validate a high-throughput system for FRET-based protein characterization and to demonstrate its use in screening a library of protein-ruler constructs. This system facilitates the high-throughput capture of complete emission spectra with excellent spectral and spatial resolution, enabling precise analysis of fluorophore properties at the nanoliter-cell-culture ensemble level. Building on our prior investigations into the impact of structural protein aggregation on biomanufacturability,^15^ we now illustrate how this platform can effectively screen protein conformation via FRET-based assays. We confirmed HyCAP’s effectiveness with fluorescent beads and genetically encoded fusion proteins (see Figures 1a and 1b), demonstrating its applicability in both controlled and biologically relevant environments. This strategy paves the way for novel high-content, high-throughput screening applications in synthetic biology, protein engineering, and cellular phenotyping.

**Figure 1.**
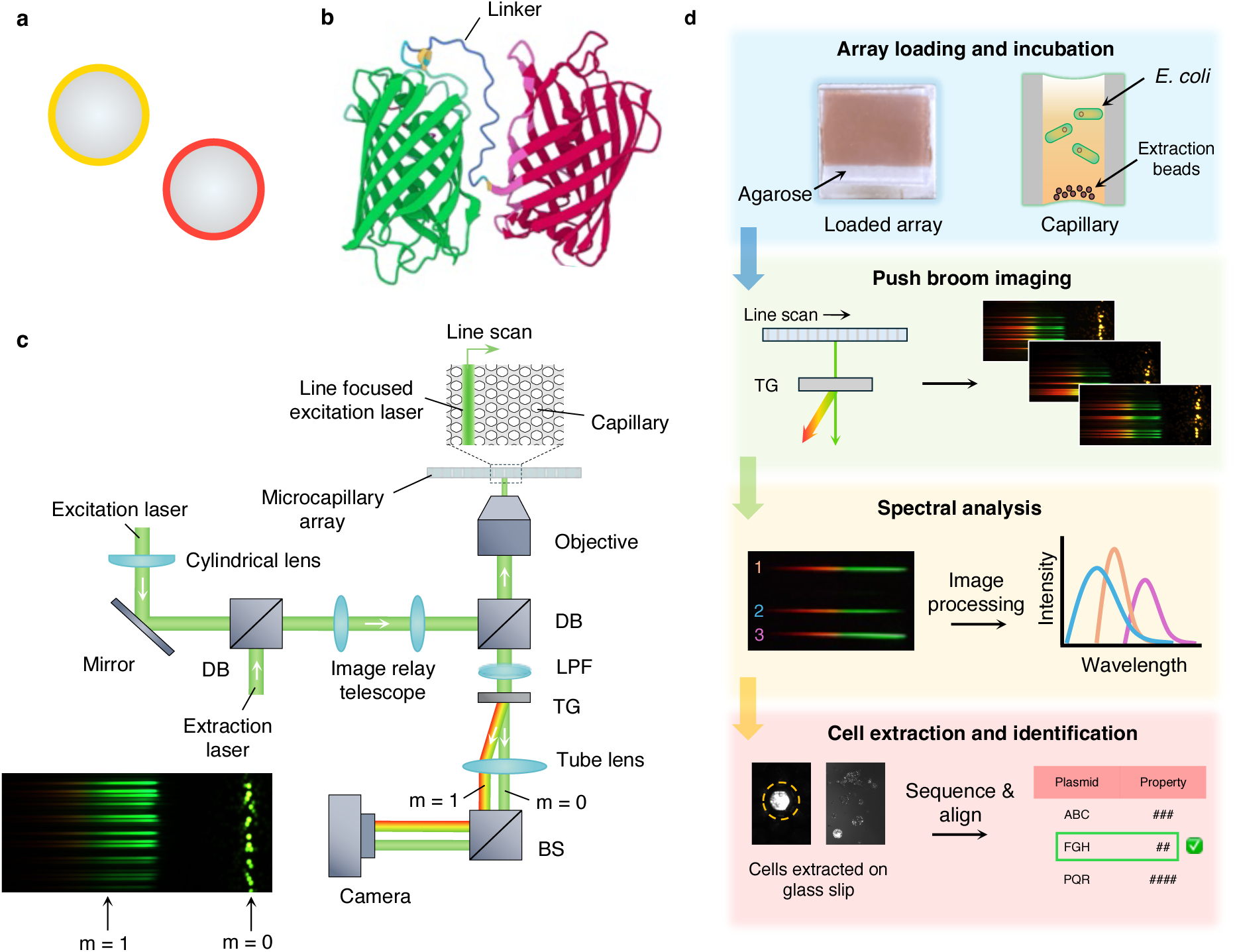
HyCAP platform and methodology. This study demonstrates screening of **(a)** fluorescent beads, and **(b)** fusion proteins. **c.** The optical pathway of the HyCAP platform is shown. White arrows show the direction of the flow of light. The snapshot illustrates a column of capillaries being illuminated by line-focused light (zeroth-order diffraction) and a dispersion band from each of the capillaries (first-order diffraction). DB: dichroic beamsplitter, LPF: long pass filter, TG: transmission grating, BS: beamsplitter. ‘m’ denotes the order of diffraction. **d.** The methodology of experiments includes loading an array with cell culture (or beads), line scanning and imaging after incubation, image processing to recover the spectral information, and recovery of capillary contents.

## RESULTS

### Hyperspectral Microscopy Platform

We used microcapillary arrays as substrates to create spatially separated miniature cultures. The platform features a 10^5^-10^8^ grid of 20-micrometer-sized glass capillaries into which cells are loaded, so that, on average, each capillary contains only one cell. ^16^ The cells grow into isogenic cultures during incubation, expressing fluorescent biomarkers for detection and identification. The array is imaged with an inverted fluorescence microscope in an automated raster scan. Bright field and fluorescence intensity are measured within each capillary, enabling the creation of a distribution of cellular properties encoded in the optical signal.

To enable hyperspectral imaging, we made two significant improvements compared to our earlier work.^15^ First, we improved the optical path by placing a cylindrical lens downstream of the collimated excitation laser (Figure 1c). This creates a line-focused beam with a width similar to a capillary’s diameter, thus exciting a column of capillaries instead of using full-field excitation. Second, we incorporated a transmission grating into the emission optical path to disperse the emitted light from each capillary within the corresponding excitation column. This setup produces an image that includes both the zeroth-order diffraction band, which corresponds to the undispersed capillaries, and the first-order diffraction band, which shows the spectrally dispersed emission components (as shown in the inset, Figure 1c). HyCAP combines the core concepts of microscopy and spectroscopy analysis with hyperspectral screening of spatially isolated clones.

### Linear dispersion characteristics

We employed a spectrophotometer coupled with an optical fiber to characterize the linear dispersion metrics of HyCAP (Figure 2a). Light beams at wavelengths ranging from 530 nm to 640 nm were transmitted through 20 µm capillaries, and the corresponding positions of these wavelengths in the first-order diffraction band were recorded (Figure 2b). Analysis of these positions revealed a linear dispersion of about 14 pixels per nanometer, with a consistent linear trend across the measured range (Figure 2c). This linearity ensures that the shape of the recorded emission spectra of fluorophores is not distorted in the process. Furthermore, a theoretical estimate of the linear dispersion—derived from the diffraction equation, optical tube length, and camera pixel size (as detailed in the Supplementary Information)—was found to be in close agreement with the experimentally measured value.

**Figure 2.**
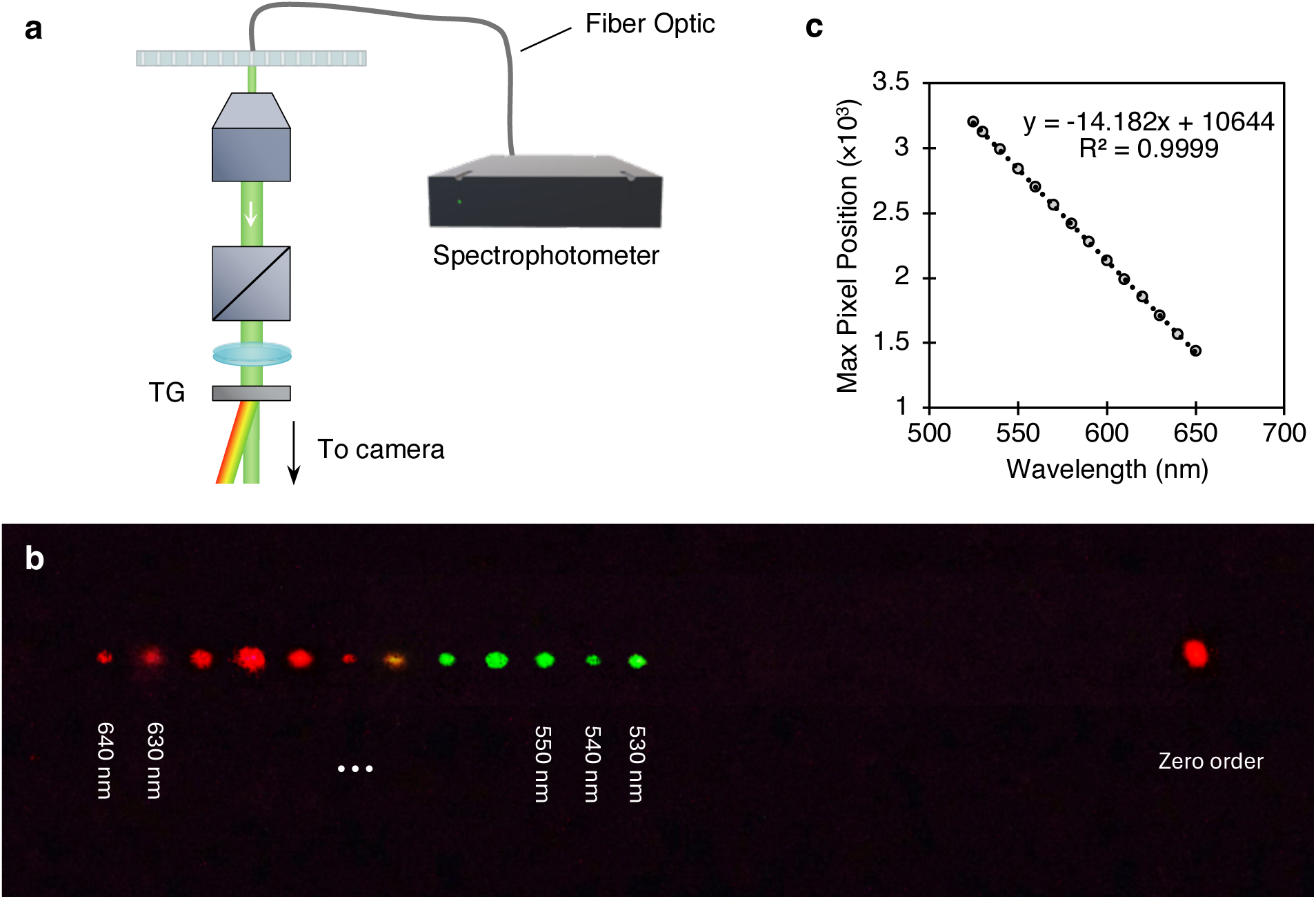
Linear dispersion metrics. **a.** For calibration, a fiber optic-coupled spectrophotometer was used to transmit narrowband wavelength light through the capillaries. **b.** The zeroth and first-order diffraction were recorded for wavelengths ranging from 530 nm to 640 nm. **c.** The dispersion relation was confirmed to be linear, with a resolution of about 14 pixels per nm.

### Fluorescent Beads Screening

To initially demonstrate the platform’s ability for spectral analysis, we used Yellow and Nile Red fluorescent beads. A representative frame (2880×1800, Figure 3a) shows the distinct first-order diffraction bands with nearly no red pixels for Yellow fluorescent beads (see full playback in the Supplementary Video). The walls of the capillaries, with a pitch of approximately 25 µm, ensure the separation of bands emanating from neighboring capillaries. We recorded the spectra (Figures 3b and 3c) of the beads across 1,050 wavelength channels (pixel indices from 600 to 1650), enabling the collection of continuous spectral profiles, unlike the 10 discrete channels used in conventional multispectral imaging. This high-resolution spectral data allows for accurate characterization of the distribution of spectral shapes and peak intensities among the beads. Furthermore, the spectra of the beads obtained from HyCAP show excellent agreement with those measured using a monochromator (Figure S1).

**Figure 3.**
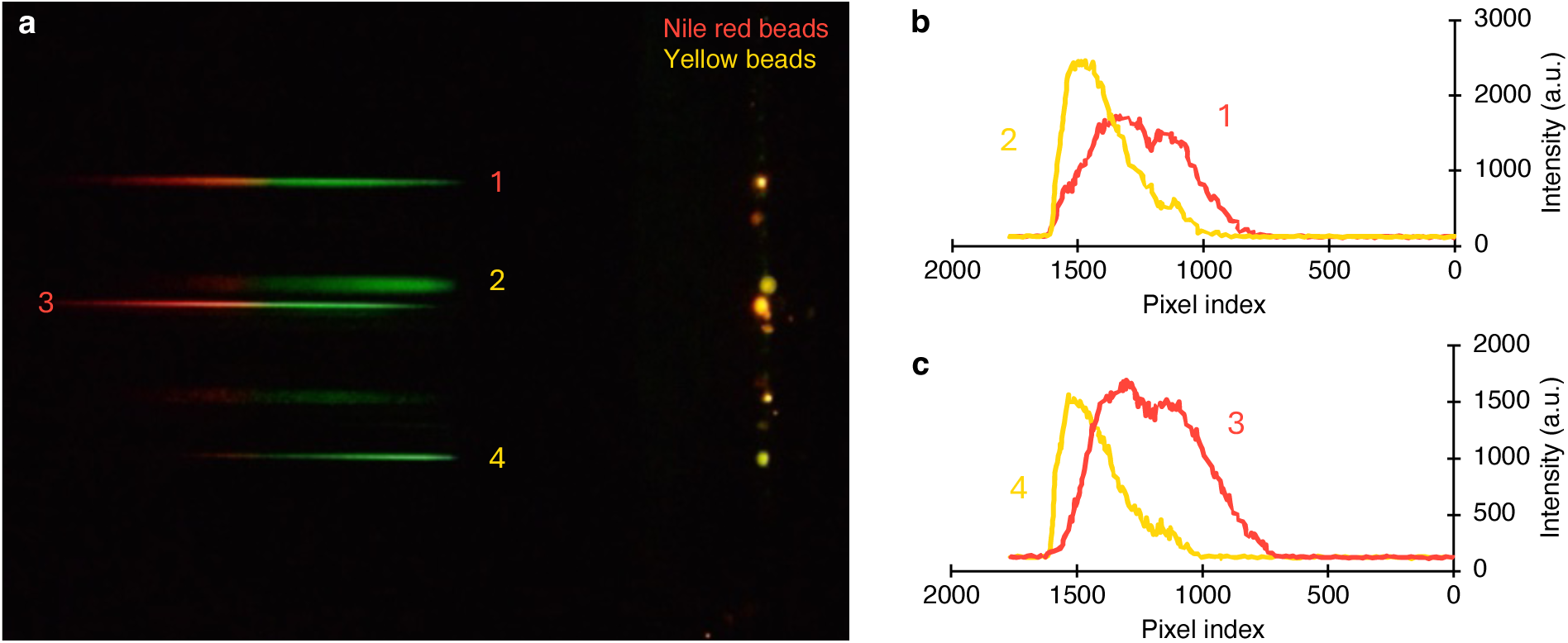
Fluorescent beads screening. **a.** Snapshot showing diffraction bands for a column of capillaries loaded with yellow and Nile red fluorescent beads. **b, c.** Representative spectra obtained for the beads are shown, capturing relative intensity as well as wavelength information.

### Ratiometric FRET

To demonstrate the suitability of HyCAP for functional screening, we chose mNG and mSc proteins as a FRET donor-acceptor pair. These fluorophores are among the brightest in their respective green and red categories, making them excellent choices for sensitive fluorescence detection measurements.^17, 18^ We constructed fusion proteins using three different linkers (Figure 4a) varying in contour length and amino acid composition.^19, 20^ The J23100 promoter was chosen (transcription rate ≈ 6×10³ au) across all plasmids, and the ribosome binding sites were designed to achieve similar translation rates (5×10⁴ au) to prevent variation in expression levels caused by DNA design (all sequences are detailed in Supplementary Information). Representative images of E. coli cultures expressing all plasmid constructs are shown in Figure 4b.

**Figure 4.**
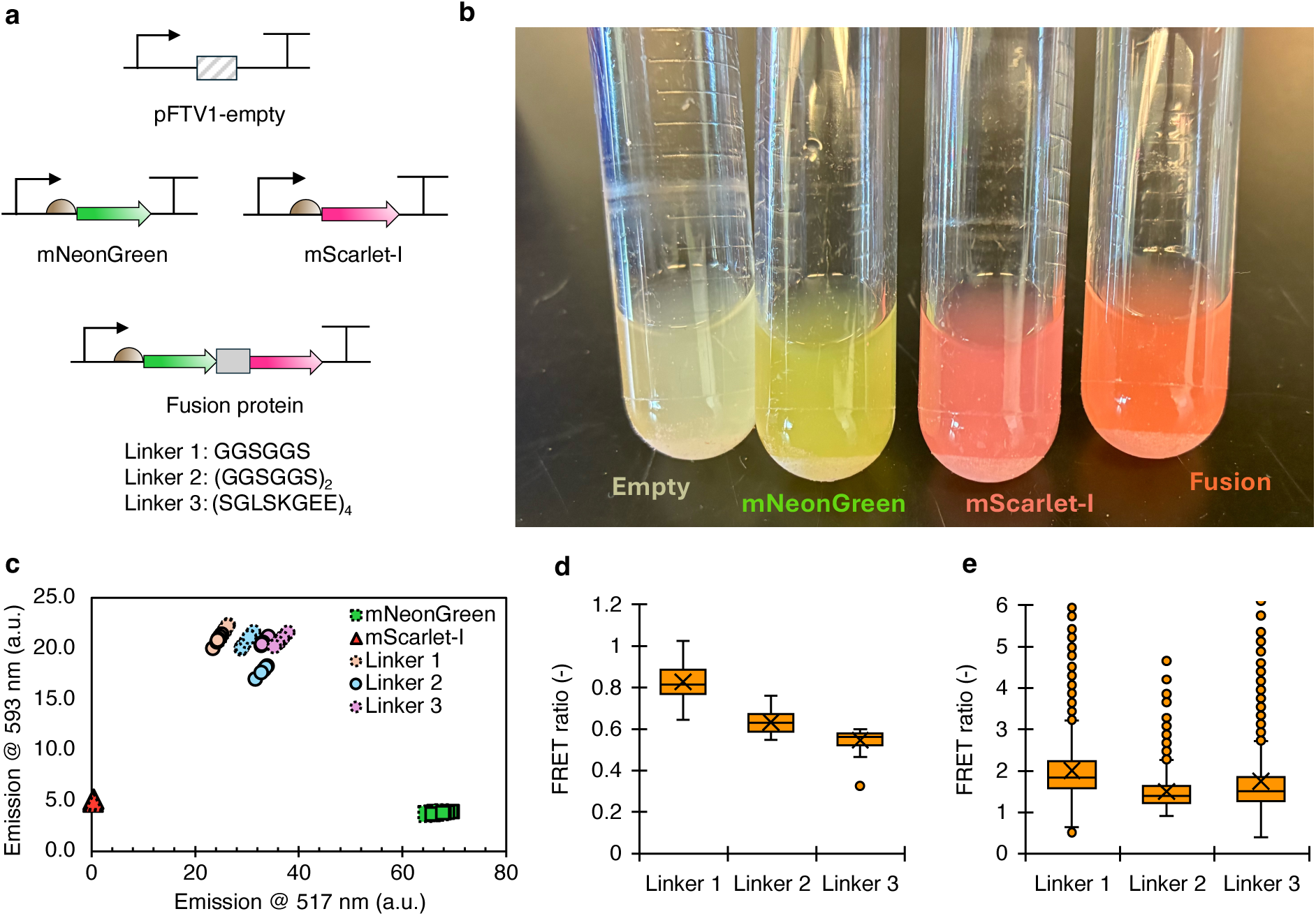
FRET constructs and ratio. **a.** Plasmid designs of various constructs are illustrated. The sequences of three linkers used in the study are shown. **b.** Representative image of *E. coli* cultures expressing each construct. **c.** Scatter plot of fluorescence intensity obtained in the green and red channels with plate reader. The dotted and solid circles denote two biological replicates per sample with eight technical replicates each. FRET ratios measured by HyCAP are significantly higher than those determined using the plate reader, which results from differences in the excitation or emission measurement setups. FRET ratio distributions of replicates measured using plate reader (30 replicates each) **(d)** and HyCAP **(e)**.

**Figure 5.**
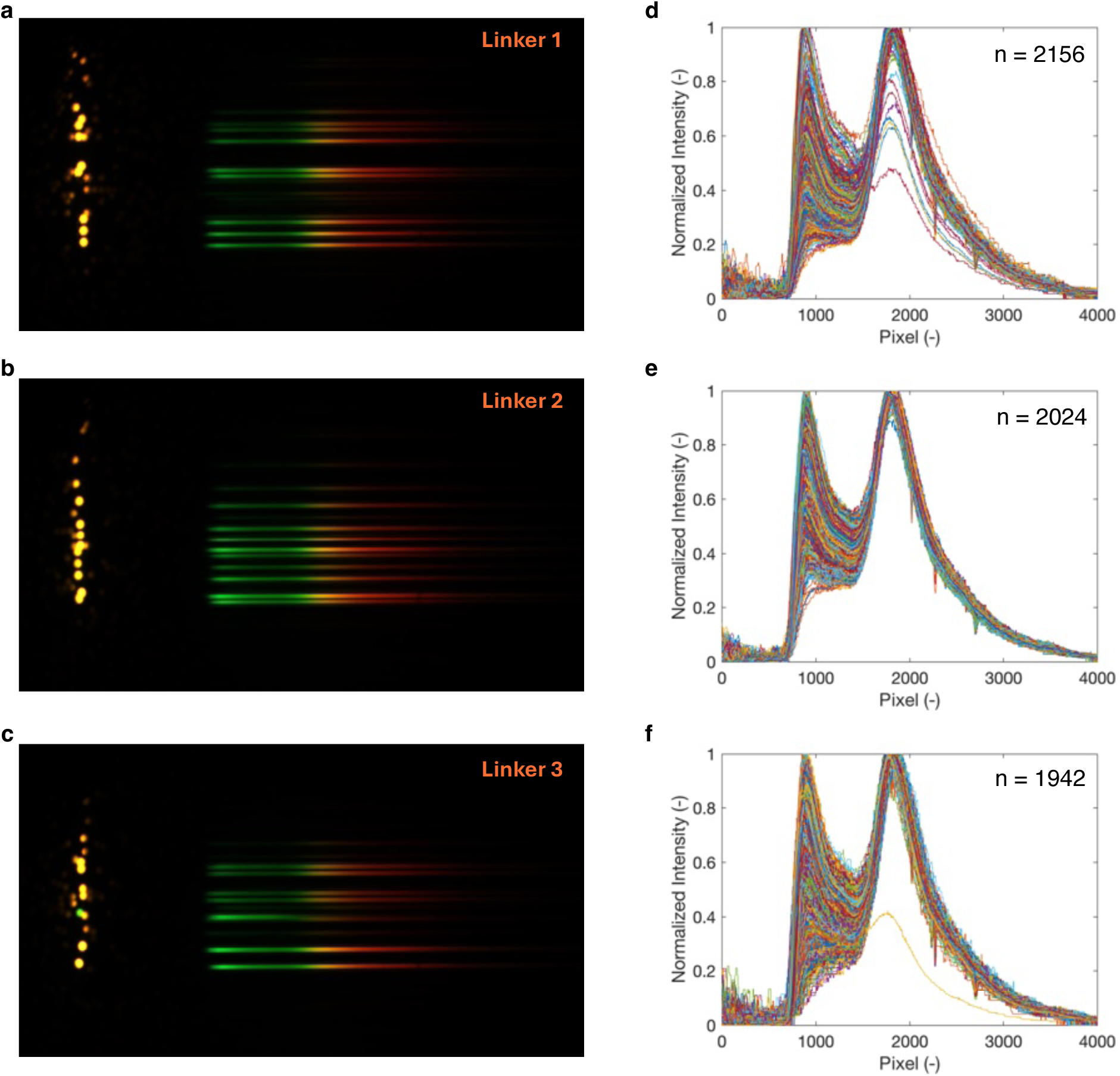
Fusion protein hyperspectral imaging. Representative snapshots recorded for Linker 1 **(a)**, Linker 2 **(b)**, and Linker 3 **(c)**. Raw spectra (‘n’ denotes frequency) of Linker 1 **(d)**, Linker 2 **(e)**, and Linker 3 **(f)**.

We initially confirmed FRET occurrence in *E. coli* expressing these constructs through fluorescence measurements from a plate reader. A scatter plot (Figure 4c) shows the raw emission intensities for all constructs in the channels tuned to peak emissions for mNG (517 nm) and mSc (593 nm) proteins, excited at 490 nm. As expected, cells expressing only mNG emitted strongly in the green channel and weakly in the red channel, consistent with its known emission spectrum (Figure S2). For the MSc protein, emission intensities in both green and red channels were low due to minimal absorbance at 490 nm. Unlike single fluorophores, cells expressing the fusion proteins showed a marked shift in emission: the red channel signal increased more than 4 times, while green channel intensity decreased by about half, indicating FRET-mediated depletion of mNG by mSc.

While measurements within replicates for each linker showed minimal variance, we observed significant variability between replicates (Figure 4c). This inter-replicate variation is probably due to differences in the stoichiometry of protein expression levels among cultures, which we link to the cytotoxicity from mSc overexpression. Whole plasmid sequencing during the cloning of mSc at 37°C revealed recombination with the E. coli host genome within the protein coding sequences. This led to a bimodal distribution of plasmid lengths, with a higher frequency of mutated plasmids compared to the designed ones (Figure S3). No such mutations were seen when cloning mNG.

We suggest that similar mutations in fusion proteins were a response to the expression burden of mSc. We observe that most mutations during cloning of fusions aimed to disrupt mSc, but not mNG. Based on this, we performed cloning and culture experiments at 30°C and 25°C, respectively, to reduce recombination and preserve the original plasmid design. Whole plasmid sequencing histograms of the purified plasmids from the modified protocol confirmed that the original designs were conserved in cultures, with the original design as the consensus sequence (Figure S4). Although recombination of the mSc coding region was minimized, it cannot be completely ruled out. We verified the stoichiometric differences in mNG and mSc expression using our microcapillary array platform, which employed full-field illumination and acceptor photobleaching (Figure S5). The photobleaching helped explain the non-FRET fluorescence.

We computed the FRET ratio through sensitized emission to provide a qualitative measure of FRET efficiency ^21–24^:

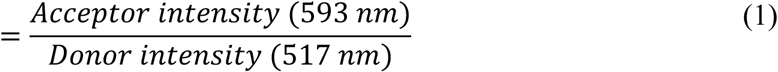

We compared the variance in FRET ratio measurements obtained using the plate reader with those from HyCAP to demonstrate the platform’s throughput and scale (Figures 4d and e, and Figure S6). About 2×10³ individual spectra (raw spectra shown in Figure S7) were collected for all fusion constructs in under 5 minutes, and the FRET ratio was calculated using the same bandwidth across the peak intensities of mNG and mSc as with the plate reader. HyCAP enabled a more robust quantification of intra-variant variability in stoichiometry due to its superior throughput and sensitivity, recording an average coefficient of variation four times greater. Additionally, the larger sampling size with HyCAP revealed a somewhat higher mean FRET ratio for Linker 3. The platform also detected several capillaries—out of thousands of replicates—primarily expressing mNG, which are rare cases of extreme stoichiometric differences in expression (Figure S8). This improved resolution allowed us to identify subtle differences in expression and FRET across the linkers with greater statistical power, thus showcasing HyCAP’s potential for high-content functional screening at much smaller cellular populations.

A wide range of mNG signal depletion was observed across capillaries, which directly affected the calculated FRET ratios. While the plate reader averaged signals across bulk populations, masking cell-to-cell differences, HyCAP captured this heterogeneity with significantly higher accuracy by reducing the cell ensemble size by over 1,000 times—from 100 µl to 0.3 nl. The signal-to-noise ratio (SNR) provided by any optical system becomes increasingly important for quantification as cellular ensemble size decreases. The background fluorescence intensity with the platform was low, allowing identification of dispersion bands with SNRs consistently over 10 in all snapshots (discussed in detail in Figures S9 and S10). Having multiple capillaries illuminated in each column per snapshot enabled us to select dispersion bands with SNR > 20 for data analysis among capillaries with a range of expression levels. As a result, the hyperspectral measurements showed a broader and more informative distribution of FRET ratios, supporting a more detailed understanding of genotype-to-phenotype connections.

### Linear unmixing and FRET efficiency

While ratiometric FRET is a common method for estimating FRET efficiency, it is inherently limited in accuracy because it is influenced by factors like donor bleed-through (i.e., fluorescence emission passing into an incorrect detection channel due to spectral overlap), cross-talk (multiple fluorochromes are excited with the same wavelength because their excitation spectra partially overlap), and spectral changes. Also, using narrow wavelength bands is not ideal for measuring fluorescence intensities in multi-fluorophore assays, as it ignores the characteristic shape of fluorophore emissions. Linear unmixing provides a more accurate and reliable method by separating each fluorophore’s contribution, enabling more precise quantification of their relative amounts. Using reference spectra for this purpose corrects these errors.

We performed high-throughput linear unmixing of FRET spectra using HyCAP. To do this, we first collected raw emission spectra from cells expressing only the individual fluorophores—mNG and mSc—and averaged them to create representative reference spectra (Figure 6). These reference spectra were then used to fit the FRET spectra obtained from fusion constructs (Figure 7a), allowing us to calculate FRET efficiency, *E*:

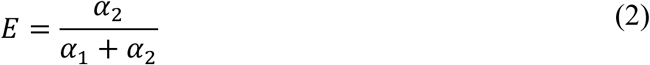

where, α_1_ and α_2_ are the relative abundances of mNG and mSc, obtained from linear least squares fitting of the respective emission spectra I(λ), per the following equation:

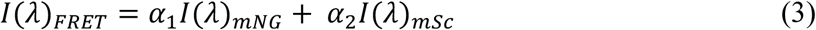

where λ denotes the wavelength.

**Figure 6.**
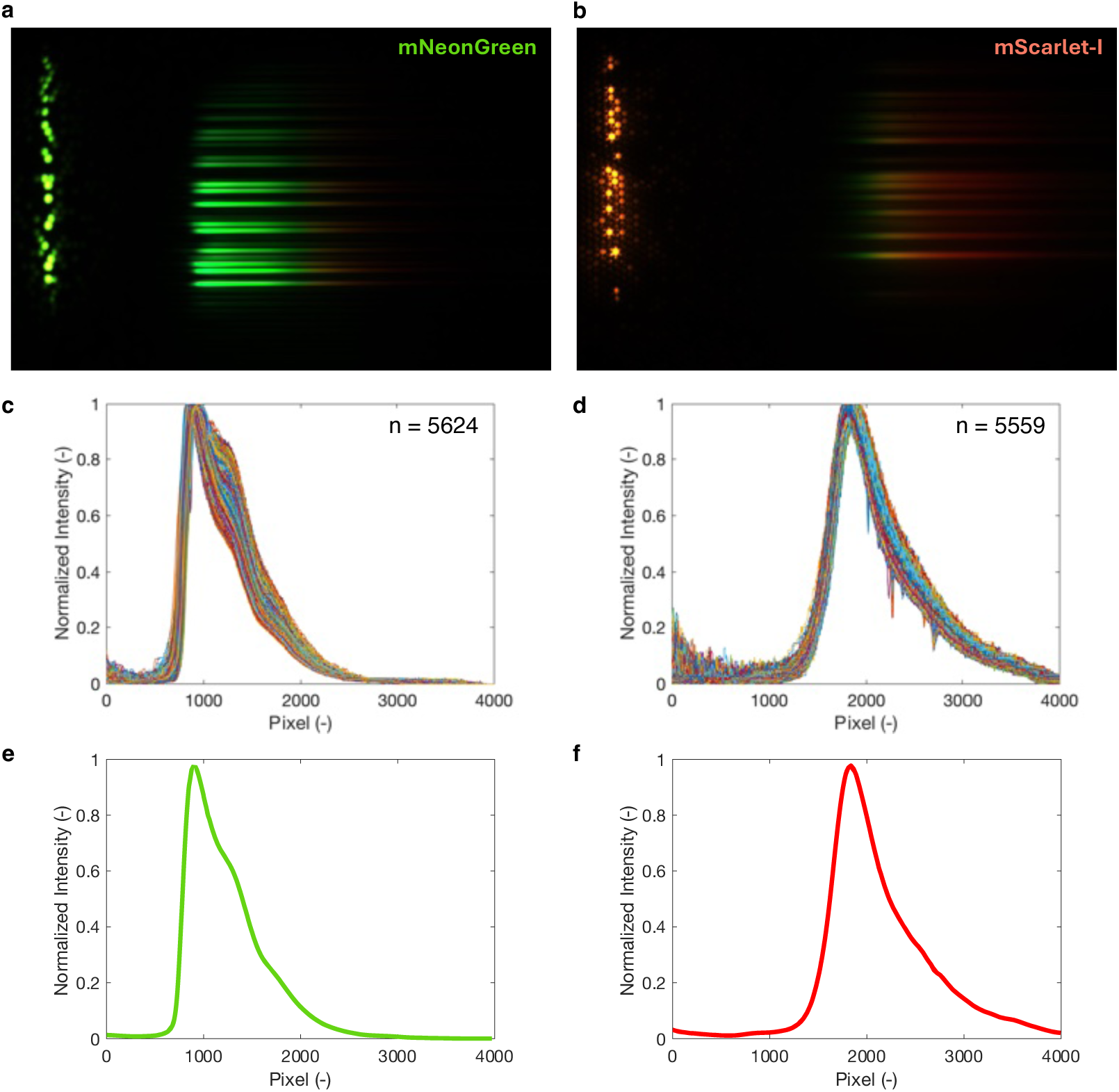
Single fluorophore hyperspectral imaging. Representative snapshots recorded for mNG **(a)** and mSc **(b)**. Raw spectra (‘n’ denotes number of microcapillaries analyzed) of mNG **(c)** and mSc **(d)**. Averaged spectra of mNG **(e)** and mSc **(f)**.

**Figure 7.**
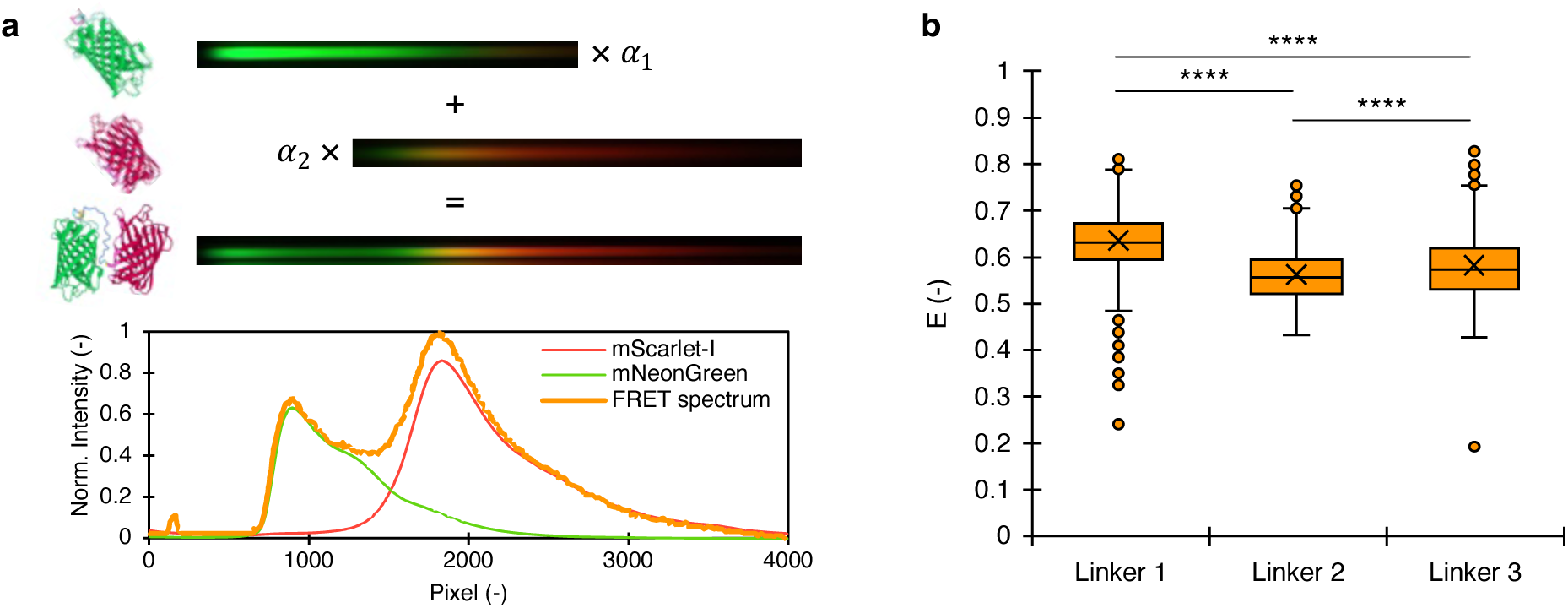
Linear unmixing and FRET efficiency. **a.** Plot depicting linear unmixing of a representative FRET spectrum using averaged mNG and mSc spectra. **b.** FRET efficiency (E) of all the fusion protein variants. Cross denotes mean value.

Notably, we observed deviations in the mNG emission profile compared to those reported in the literature. ^25, 26^ Specifically, a relative shift in intensity was detected near the 1330-pixel location (∼545 nm), manifesting as a shoulder in the averaged FRET spectrum. The variation in the relative intensity of this shoulder indicates that it is intrinsic to the mNG structure rather than caused by the instrument. The absence of a similar shoulder in the mSc spectra further supports this. We conducted an experiment with significantly lower ensemble levels (after 12 hours of incubation compared to 48 hours) to isolate the potential effect of protein aggregation within cells on mNG emission (Figure S11). A 35-fold increase in exposure was needed to record the spectra at the shorter incubation period (yet with lower peak intensities), confirming poorer expression of mNG in capillaries. However, the shoulder artifact around the 1330-pixel location was still observed. A population of mNG molecules with red-shifted emission appears to be present. This suggests that the artifact in mNG emission is influenced by the E. coli cellular environment, possibly due to misfolding or interactions rather than aggregation. In contrast, the emission profile of mSc remained consistent, with no significant changes in spectral shape. These findings demonstrate the power of our platform in resolving complex and unexpected spectral features, which are essential for accurately quantifying FRET efficiency.

The FRET efficiency trend depicted in Figure 7b is highly significant (*p* < 1^-10^) for all linker combinations, indicating a strong statistical basis for relative loss or gain in FRET efficiency. Moreover, the effect sizes (A statistical measure that indicates whether the difference between the means of two data sets is statistically significant) of Linkers 1& 2 and Linkers 1&3 are large (> 0.8) according to Cohen’s d test (Table S1). Linker 2 was designed with a contour length twice that of Linker 1. These Gly-Ser flexible linkers have been modeled using the worm-like chain (WLC) model,^27^ in which the end-to-end distance is a function of both the contour length and the persistence length (found to be 4.5 Å compared to 3.8 Å distance between adjacent *C*_α_ atoms).

According to this model, the increased contour length of Linker 2 results in a greater average end-to-end distance, thereby reducing the efficiency of FRET compared to Linker 1—a trend that was observed experimentally in literature as well as here.

The FRET efficiency trend shown in Figure 7b is highly significant (*p* < 1^-10^) for all linker combinations, indicating a strong statistical basis for the relative reduction or increase in FRET efficiency. Additionally, the effect sizes of Linkers 1&2 and Linkers 1 &3 are large (> 0.8) according to Cohen’s d test (Table S1). Linker 2 was designed with a contour length twice that of Linker 1. These Gly-Ser flexible linkers have been modeled using the worm-like chain (WLC) model, ^27^ in which the end-to-end distance depends on both the contour length and the persistence length (found to be 4.5 Å compared to 3.8 Å, the distance between adjacent *C*_α_ atoms). According to this model, the increased contour length of Linker 2 results in a greater average end-to-end distance, thus reducing FRET efficiency compared to Linker 1—a trend observed both in literature and in our experiments.

Although Linker 3 has a contour length more than twice that of Linker 2, it shows a FRET efficiency similar to that of Linker 2, indicating a shorter end-to-end distance due to a more compact structure. We used computational modeling to explore its possible conformations since the structure of Linker 3 has not been studied experimentally before. Simulated models generated with PEP-FOLD4, I-TASSER, and AlphaFold3 (Figure S12) consistently predicted a helical shape, specifically a folded or kinked helix likely caused by charged amino acids in the sequence. Such a conformation could effectively reduce Linker 3’s end-to-end distance despite its increased contour length, leading to a FRET efficiency similar to Linker 2. These results suggest that, beyond length, the secondary structure of the linker is crucial in determining the proximity of donor and acceptor fluorophores and thus energy transfer efficiency. While further studies are needed to confirm Linker 3’s conformation, which is outside the scope of this discussion, our work highlights the potential of HyCAP in detecting protein conformational changes.

## DISCUSSION

We have developed a hyperspectral microscopy system that provides high-throughput imaging of up to 15 clones per second (1 image snapshot per second) and high-resolution spectral analysis (e.g., This rate can be increased by 10 to 100 times with faster electronics and improved software processing). The platform uses a transmission grating to disperse wavelength components in fluorescence emission from fluorophores across 103 channels. The HyCAP imaging modality is automated and requires no user intervention after setting the initial test conditions. Standard microscopy tools and software were used to create a cost-effective platform that enables highly sensitive (SNR > 10) fluorescence quantification with miniature cell cultures. Although we focused on the widely used green-red (520–700 nm) spectral range, HyCAP is suitable for fluorophores across the entire visible spectrum.

We demonstrated the capabilities of HyCAP for fluorophore engineering by measuring inter-bead variation in fluorescence, specifically in wavelength and intensity signals. The continuous spectra obtained for the fluorophores matched those from a standard monochromator-based spectrometer. HyCAP’s high throughput—thanks to microcapillary arrays and push-broom imaging—allows users to screen thousands of clones and isolate a single variant with precise laser extraction.^28^ We collected approximately 5×10^3^ spectra for both mNG and mSc, revealing notable changes in the mNG spectrum. The ability to detect such alterations—and their magnitude—is essential in fluorophore engineering (and FRET), and such information would likely be lost with multispectral techniques due to their low spectral resolution, typically around 10 wavelength channels.

We used the cytotoxicity of mSc to introduce stoichiometric variation in fusion protein expression levels and to simulate screening of libraries of FRET clones. This variability was first confirmed with plate reader measurements. However, while plate readers are helpful for bulk fluorescence measurement, they are inherently low-throughput and not ideal for screening large numbers of clones needed to capture the full range of expression differences. Additionally, plate readers usually measure fluorescence within narrow spectral ranges, which limits their accuracy for FRET, as corrections for donor bleed-through and environmental spectral changes are needed (as seen with mNG in our case). Our platform overcomes these limitations by enabling: 1. screening of larger clone-to-clone heterogeneity with ensembles 10^3^ times smaller at high throughput, and 2. linear unmixing of individual fluorophore amounts and correction for donor bleed-through. Moreover, the ensemble size can be adjusted by controlling the incubation time.

It has been demonstrated that linear unmixing yields consistent FRET estimates, unlike other common methods such as one-filter set, two-filter set, three-filter set, and peak intensities.^29^ Although we do not correct for direct acceptor excitation in our studies due to the minimal contribution from mSc excitation at 505 nm (absorbance ∼ 0.2), this correction can be easily implemented on the platform by using an excitation laser at which donor absorbance is zero, or by performing another line-scan to record acceptor emission when the acceptor is excited, although this may reduce throughput. For a platform mainly designed for high-throughput sorting and enrichment of clones in libraries, ensuring consistent FRET trends with respect to end-to-end distances is essential and sufficient.

Linker proteins are used in FRET-based studies as protein rulers, such as proteins engineered with donor and acceptor fluorophores at specific distances. Linker peptides play a vital role in proteins by providing flexibility, stability, and functionality. For example, they can: (i) regulate conformation by influencing domain orientation, which affects function and interactions; (ii) facilitate protein interactions by offering binding sites or accessibility; (iii) modify mechanical properties like elasticity or rigidity. Without an appropriate linker between protein domains, there is a risk of amyloid formation and aggregation due to structural rigidity caused by decreased flexibility, increased tendency for beta-sheet formation that promotes aggregation, and reduced steric hindrance, which allows for closer interactions.

The high throughput of our platform enabled us to identify a statistically significant trend in FRET efficiency versus linker type, revealing the conformation of the linkers despite sensitivity to stochastic variance in the stoichiometry of mNG and mSc. It has been demonstrated elsewhere that high throughput is crucial when establishing genotype-to-phenotype correlations in highly variable phenomena. The decrease in FRET efficiency caused by the increased length of flexible Gly-Ser linkers was comparable to previous reports.^19^ Additionally, the platform indicated a more compact structure for Linker 3 compared to the shorter Linker 2. This finding was confirmed through computational modeling, which strongly suggested a kinked alpha-helical structure that offsets the increased contour length of Linker 3.

In summary, HyCAP is ideal for sorting and enriching large libraries of clones based on fluorescence assays. To the best of our knowledge, HyCAP is the first platform that offers sensitive, high-resolution hyperspectral imaging with throughput on the order of 10^5^ per array. Based on our previous demonstration,^28^ a library of genotypes can be expressed on the array and enriched based on desired levels of phenotype. The FRET protocol is suitable for studying protein-structure features that vary with differences in amino acid sequences as well as protein assembly. In addition to protein conformation (Figure 8), HyCAP shows potential for applications in fluorophore engineering such as emission changes related to environment (pH, enzymatic activity, temperature, etc.), stoichiometry differences, fluorophore orientation, and photobleaching through time-resolved experiments.

**Figure 8.**
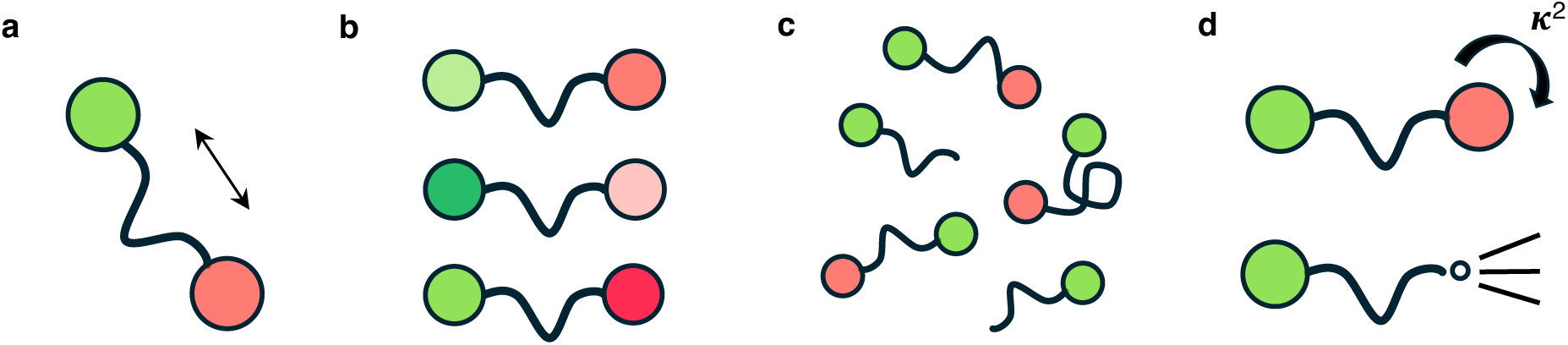
Applications of hyperspectral microscopy platform. The platform can be used to screen for conformational changes of proteins **(a)**, emission shifts **(b)**, stoichiometry differences **(c)**, fluorophore orientation, and photobleaching **(d)**.

## DATA AVAILABILITY

The authors declare that data supporting the findings of this study are available within the paper and its supplementary information files.

## AUTHOR CONTRIBUTIONS

M.C.D. conceived the project. T.M.B. and K.S. developed the platform. K.S. and B.D.A. designed the plasmids. K.S. performed the cloning, designed the experiments, and conducted the screening. K.S. wrote the manuscript together with M.C.D. All authors participated in manuscript revisions, discussion, and data interpretation.

## ACKNOWLEDGEMENT OF SPONSORSHIP STATEMENT

We thank Dr. Howard Salis for kindly providing the pFTV1-mRFP1 plasmid. We also thank Harry Adamson for assistance with the plate reader measurements.

This effort was sponsored in whole or in part by the Defensewerx. The U.S. Government is authorized to reproduce and distribute reprints for Governmental purposes, notwithstanding any copyright notation thereon.

## DISCLAIMER

The views and conclusions contained herein are those of the authors and should not be interpreted as necessarily representing the official policies or endorsements, either expressed or implied, of the Central Intelligence Agency.

## METHODS

### Instrumentation

The platform (Figure 1c) is based on an inverted fluorescence microscope (Olympus IX73). The microscope was equipped with a digital color camera (ToupTek ATR2600C), motorized XY stage (Marzhauser Wetzlar Tango 3), motorized focus drive (Marzhauser Wetzlar), and a 10× objective lens (Olympus). Custom software was designed in LabView using the parent SDKs to operate the camera, and stage. Microcapillary arrays with 20 µm capillary diameter (INCOM, Inc.) were used throughout all experiments, which contain about 8×10^5^ capillaries in total.

The details regarding the extraction laser can be found in ref^1^. For excitation, a collimated beam of light from a 505 nm laser diode (Q505-14, Qiaoba) was passed through a cylindrical lens (75 mm, LJ1703RM, Thorlabs). The linearly diverging beam was then relayed through a telescope (1×) and diverted into the rear objective aperture using a 520 nm dichroic beamsplitter (AT515DC, Chroma). The objective focused the beam into a line of width about 10 μm – 20 μm in the focal plane. The emission collected by the objective was passed through another 520 nm filter (AT515LP, Chroma) to suppress 505 nm excitation and consequently through a transmission grating (300 lines/mm, GT50-03, Thorlabs). The diffraction bands were then focused on the camera sensor (Sony IMX571) by a tube lens (180 mm).

Each pixel column within the first-order diffraction band corresponds to a distinct wavelength channel, and the APS-C (Advanced Photo System type-C) camera sensor offers wavelength channels on the order of 10^3^ for high-resolution spectral analysis. We ensure high throughput by continuous line-scanning (otherwise known as push-broom imaging ^30^, see Supplementary Video) across the array at a fast rate, capturing images at regular intervals (Figure 1d). Our camera control offers a minimum snapshot-acquisition time of about 20 ms. These snapshots are subsequently processed through image analysis software (described in detail in Supplementary Information) to reconstruct the spectral profiles. It is important to note that, while this platform is capable of laser-based recovery of specific clones, that aspect is not demonstrated here. This work focuses solely on the screening capability of HyCAP.

### Microcapillary array loading, screening, and sequencing

The arrays were sterilized before use, and treated with a plasma wand to impart hydrophilicity. Cells in Luria broth (LB) were loaded onto the array, with cell concentration adjusted such that each capillary acquired a single cell on average. After loading, the array was overlaid with an agarose gel layer (2% w/v, 1 – 2 mm thick) and incubated (at 25°C) in a sealed petri dish lined with moist wipes as desired. After screening, arrays were sterilized in 70% ethanol for several hours. Arrays were cleaned thoroughly under a DI water stream, with intermediate ultrasonication for 2 min.

### Imaging and processing

The array was placed on the sample stage, and X-coordinates of the start and end positions for the 1D scan were noted. Z-positions of the objective with the sample in focus at these X-coordinates were noted as well. According to the traverse speed of 50 μm/s along the X-axis, the traverse speed in the Z-direction was calculated to keep the array in focus throughout the scan (Figure S13).

The methodology of image processing developed (script written in MATLAB) to reveal the spectral information of emissions is illustrated in Figure S14. The color snapshot of the array was first cropped to remove the zeroth order band. The cropped image was converted to grayscale and straightened digitally by 0.25° using the ‘imrotate’ function with ‘nearest’ interpolation method. The values of all pixels in each row of the grayscale image-matrix were summed. This generated a plot with intensity peaks denoting first-order band positions along the vertical axis of the image. A minimum threshold for peak intensity was set for peak selection to obtain similar total numbers of spectra across constructs. Peak maximum indicated the band center axis, and a standard 20 pixel-rows (10 pixel-rows on either side of the center axis) were assigned as the thickness for each band. To reveal the spectrum, values of all pixels in each column (of 21 pixels) were summed and plotted.

### Plasmid synthesis and cloning

All nucleotide sequences used in this study are provided in the Supplementary Information. All gene blocks were designed on denovodna.com. The base vector for plasmid construction was pFTV1-mRFP1, from which the mRFP1 coding sequence along with its ribosome binding site (RBS) was removed by restriction digestion using BamHI-HF and PstI-HF (New England Biolabs). The resulting vector backbone was then assembled with a gene block (Genscript) containing a protein insertion site using Gibson Assembly, generating the intermediate construct pFTV1-empty.

This pFTV1-empty plasmid served as the template backbone for all subsequent plasmid constructs. Gene blocks (Genscript) containing RBSs and coding sequences for the respective fluorophores and linker-fusion constructs were assembled into pFTV1-empty using Golden Gate Assembly with (BsaI-HF v2, NEB). This process yielded the final expression constructs: pFTV1-mNG, pFTV1-mSc, and pFTV1-Linker 1 through 3 plasmids, each assembled in separate Golden Gate reactions. The plasmids were transformed into NEB Stable (recovered and plated at 30°C). The plasmids were sequence-confirmed via Oxford Nanopore whole plasmid sequencing (Quintara Biosciences). The plasmids (including pFTV1-empty) were finally transformed (recovered and plated at 30°C) into NEB BL21(DE3), and glycerol stocks of this strain were used subsequently for all experiments.

### Plate reader measurements and FRET ratio

Isogenic colonies of pFTV1-empty, pFTV1-mNG, pFTV1-mSc, and pFTV1-Linker 1 through 3 were inoculated in chloramphenicol-supplemented (35 μg/ml) LB, and incubated at 25°C and 250 RPM overnight. Endpoint measurements were carried out using a TECAN Spark plate reader at 490 nm excitation, and 517 nm and 593 nm emissions with 7.5 nm bandwidth each. The fluorescence emissions were normalized with respect to OD_600_ and background correction was done before calculating FRET ratios.

### Hyperspectral imaging, FRET ratio, and linear unmixing

pFTV1-mNG, pFTV1-mSc, and pFTV1-Linker 1 through 3 were inoculated in chloramphenicol-supplemented (35 μg/ml) LB and incubated at 25°C and 250 RPM. 100 μl aliquots of cell suspension in supplemented LB were made using 2 μl of stock culture each at OD_600_ = 0.1. Each aliquot was loaded on a separate array and incubated at 25°C for 48 hr. Line-scanning hyperspectral imaging was performed with each array with snapshots acquired at intervals of 1 s at an exposure of 10 ms each. For pFTV1-mSc, the gain of the camera was increased to 2×. To calculate single-fluorophore peak intensities using the obtained spectra, a bandwidth of 7.5 nm around the peak maximum for mNG and mSc was used. Pixel indices of peak maxima were identified using reference spectra, and a linear dispersion of about 15 pixels/nm was used to assign bandwidth. To obtain reference spectra of mNG and mSc, raw spectra were averaged. These reference spectra were normalized to the max intensity of 1 a.u. and used to deconvolute the normalized spectra obtained for each fusion protein through a custom MATLAB script employing the function ‘nlinfit’. Another set of FRET ratios was also calculated using the peak maxima identified through linear unmixing.

