## supplemental figures and tables for "High-Throughput Screening of FRET-Based Protein Rulers Using a Hyperspectral Microcapillary Array"

### Hyperspectral Microscopy Platform for High-Throughput FRET Assay Screening

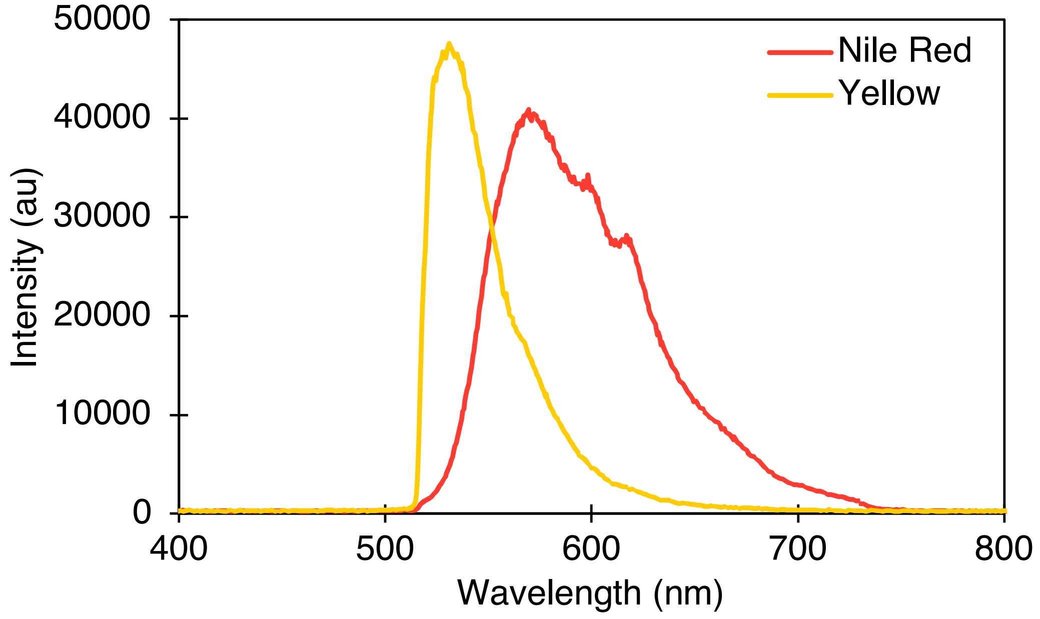


**Figure S1. Control fluorescence emission data.** Emission spectra of Yellow and Nile red fluorescent beads as obtained using Ocean Optics monochromator are shown.

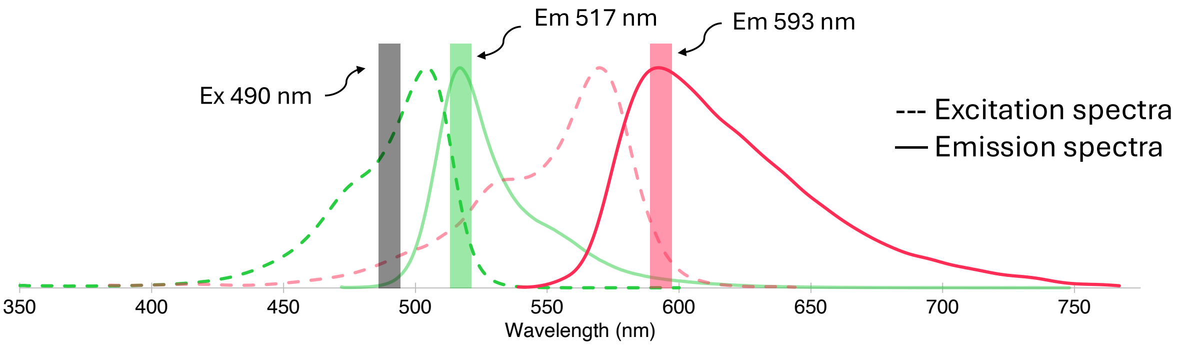


**Figure S2. Excitation and emission characteristics of FRET probes.** Excitation (dashed line) and emission (solid line) spectra of mNeonGreen (in green) and mScarlet-I (in red) are shown as obtained from fbase.org. The fusion constructs were excited at 490 nm, and emission was measured in two channels, 517 nm and 593 nm, each with a bandwidth of 7.5 nm.

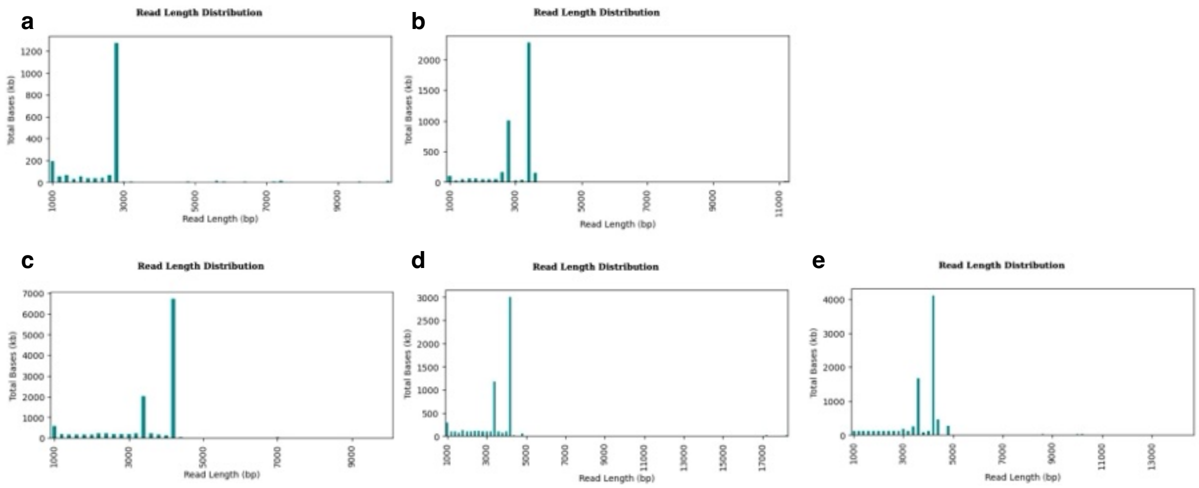


**Figure S3. Initial sequencing results for the constructs. a.** Read length of mNG plasmid is correct and equal to the designed plasmid. Read lengths of mSc (**b**), Linker 1 (**c**), Linker 2 (**d**), and Linker 3 (**e**) follow a bimodal distribution with the dominating mutated plasmid of the longer read length.

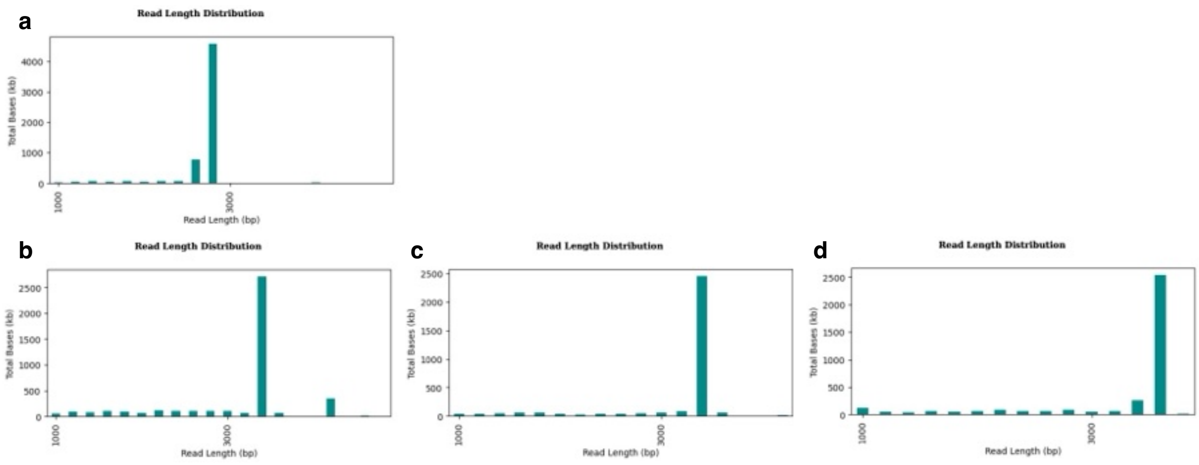


**Figure S4. Sequencing results with the modified cloning protocol.** Read lengths of mSc (**a**), Linker 1 (**b**), Linker 2 (**c**), and Linker 3 (**d**) follow a unimodal distribution with the correct plasmid length.

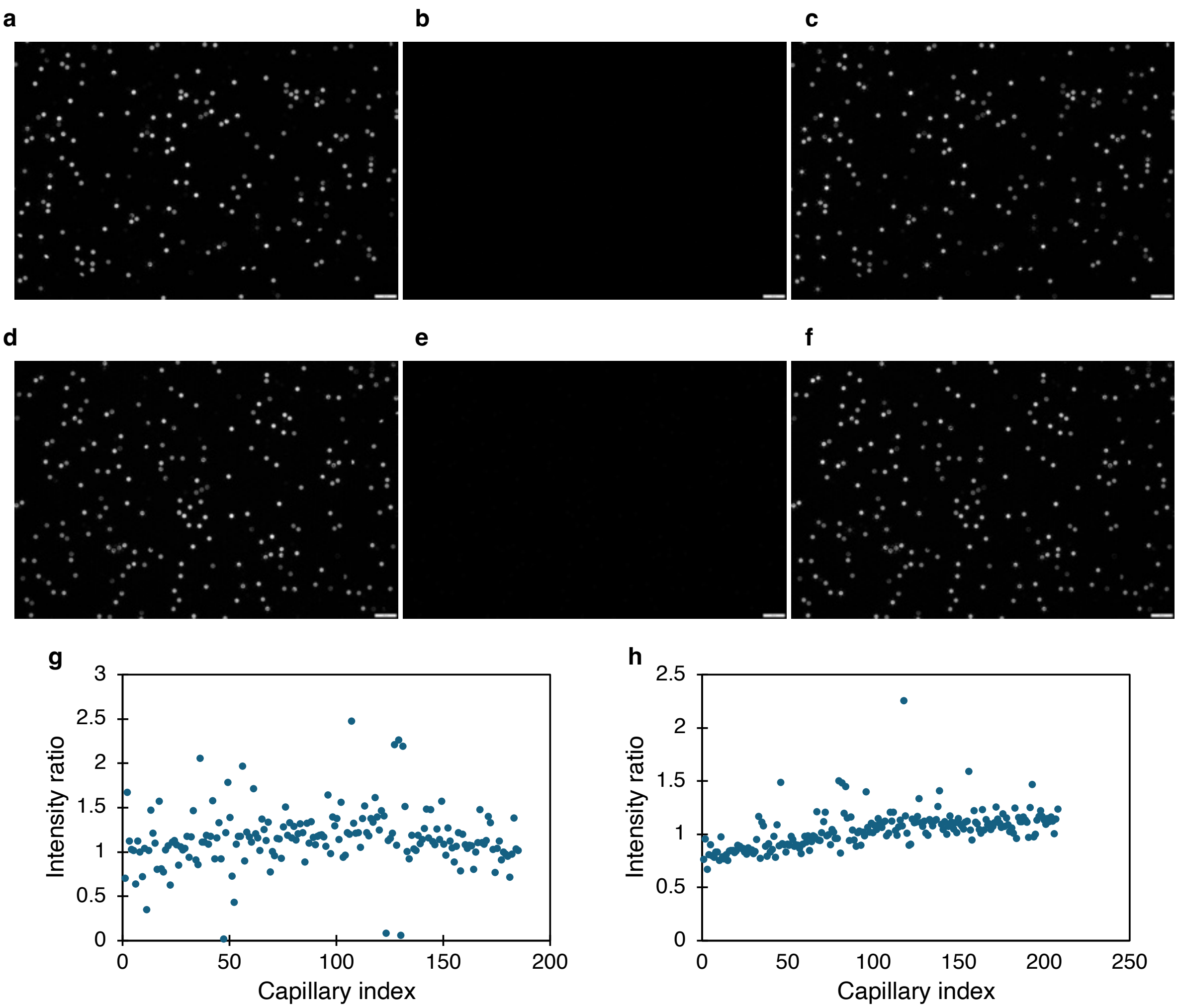


**Figure S5. Acceptor photobleaching with Linker 3.** Fluorescence images of two locations on the array of Linker 3 are shown. **a, d.** Initial fluorescence images of mSc. **b, e.** Fluorescence images of mSc recorded after photobleaching at the same exposure. **c, f.** Fluorescence image of mNG. Intensity ratios of mSc to mNG calculated for the two images (**g** for **a**, **h** for **d**).

The experiment was conducted via wide-field fluorescence illumination using the CoolLED pE400$^{max}$. Firstly, mSc fluorescence images were recorded using the 550 nm channel. Then the capillaries were continued to be illuminated with LED light until the mSc fluorophores were bleached. Then using the 450 nm channel, mNG fluorescence was recorded at the same locations. Using a custom image processing script, capillary intensities in both channels were estimated and mSc:mNG intensity ratios were calculated.

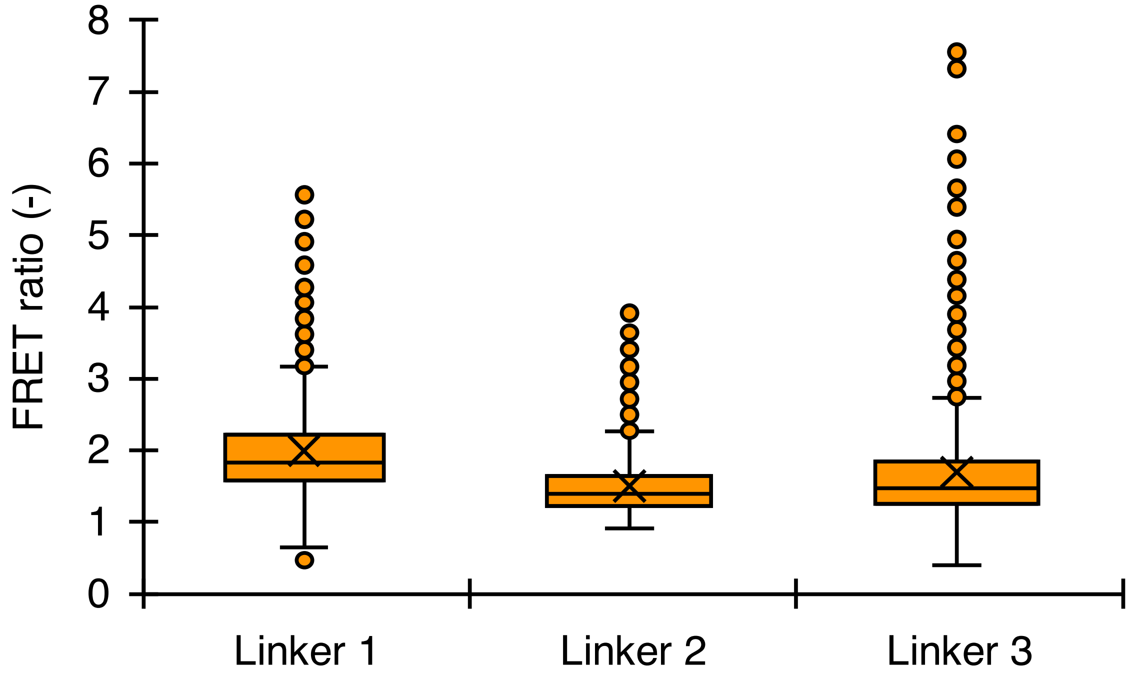


**Figure S6. FRET ratio, related to Figure 4 in main text.** FRET ratio calculated using peak intensity wavelengths corresponding to fitting parameters of linear unmixing.

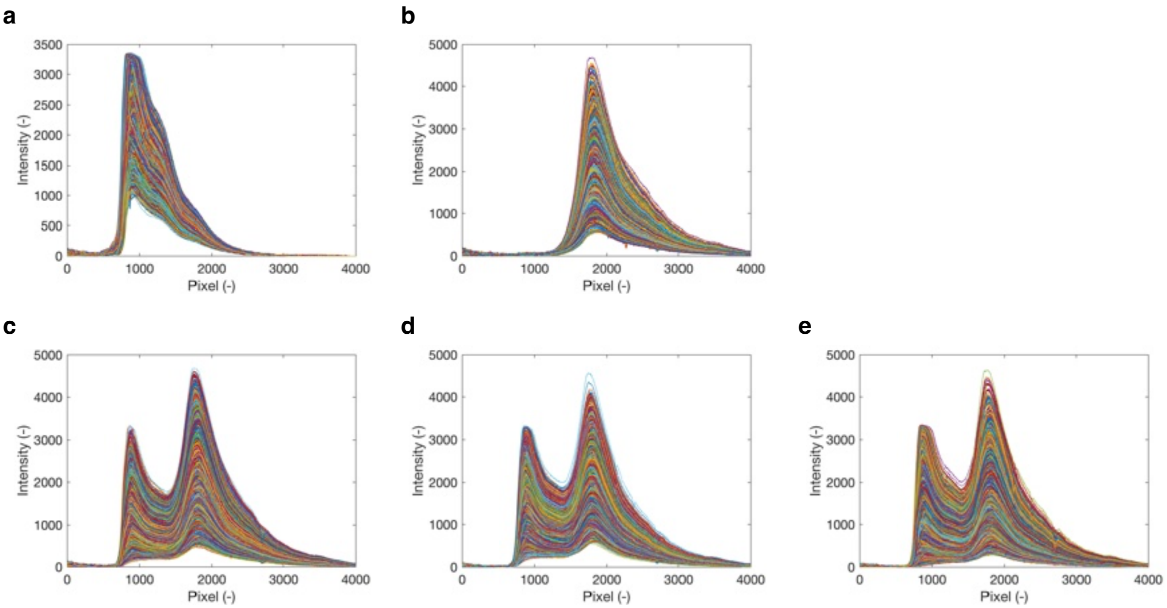


**Figure S7. Raw spectra collected with HyCAP, related to Figures 5 and 6 in main text.** Raw spectra collected using HyCAP for mNG (**a**), mSc (**b**), Linker 1 (**c**), Linker 2 (**d**), and Linker 3 (**e**) are depicted.

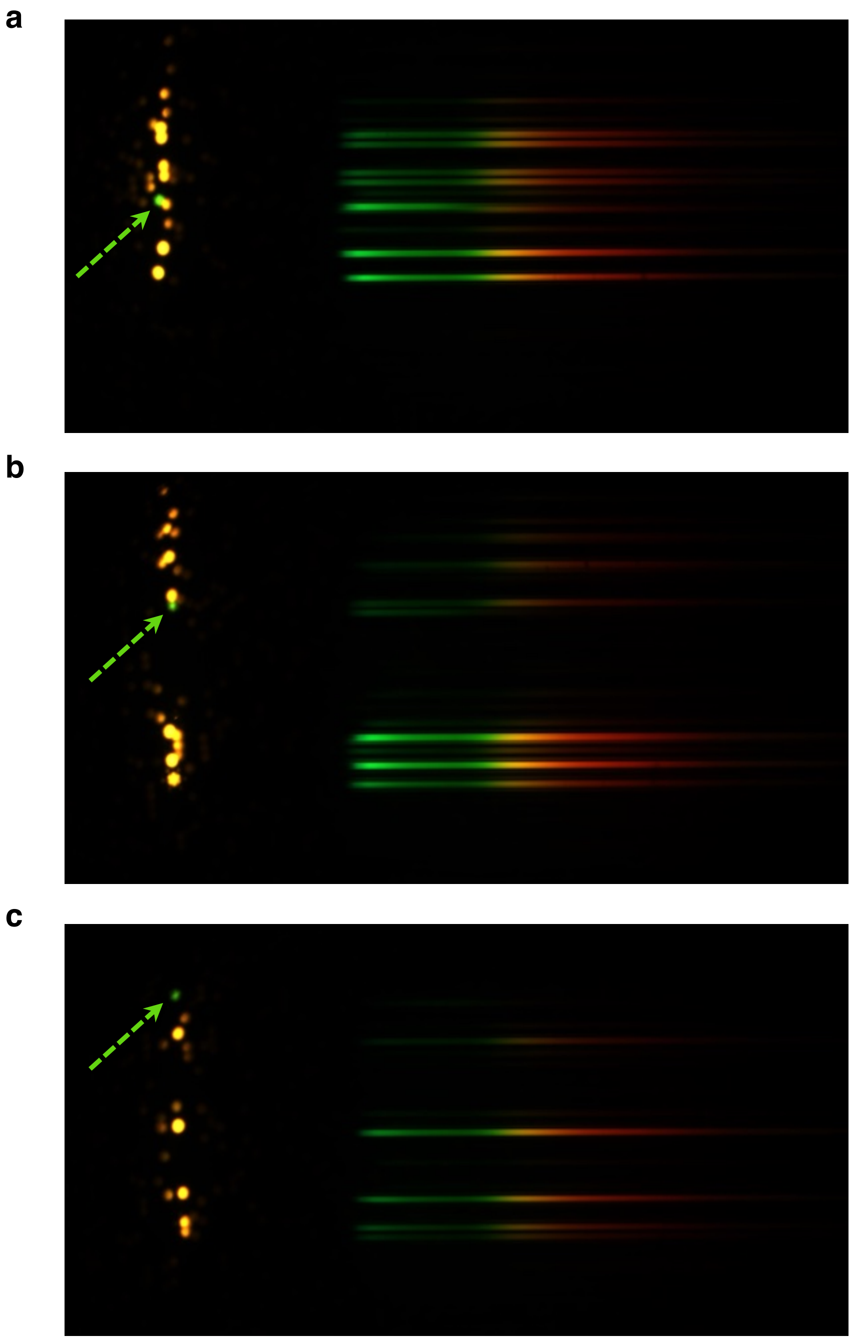


**Figure S8. Stoichiometric differences in expression.** Representative images (**a – c**) of linker constructs expressing predominantly mNG. These capillaries are indicated with green arrows.

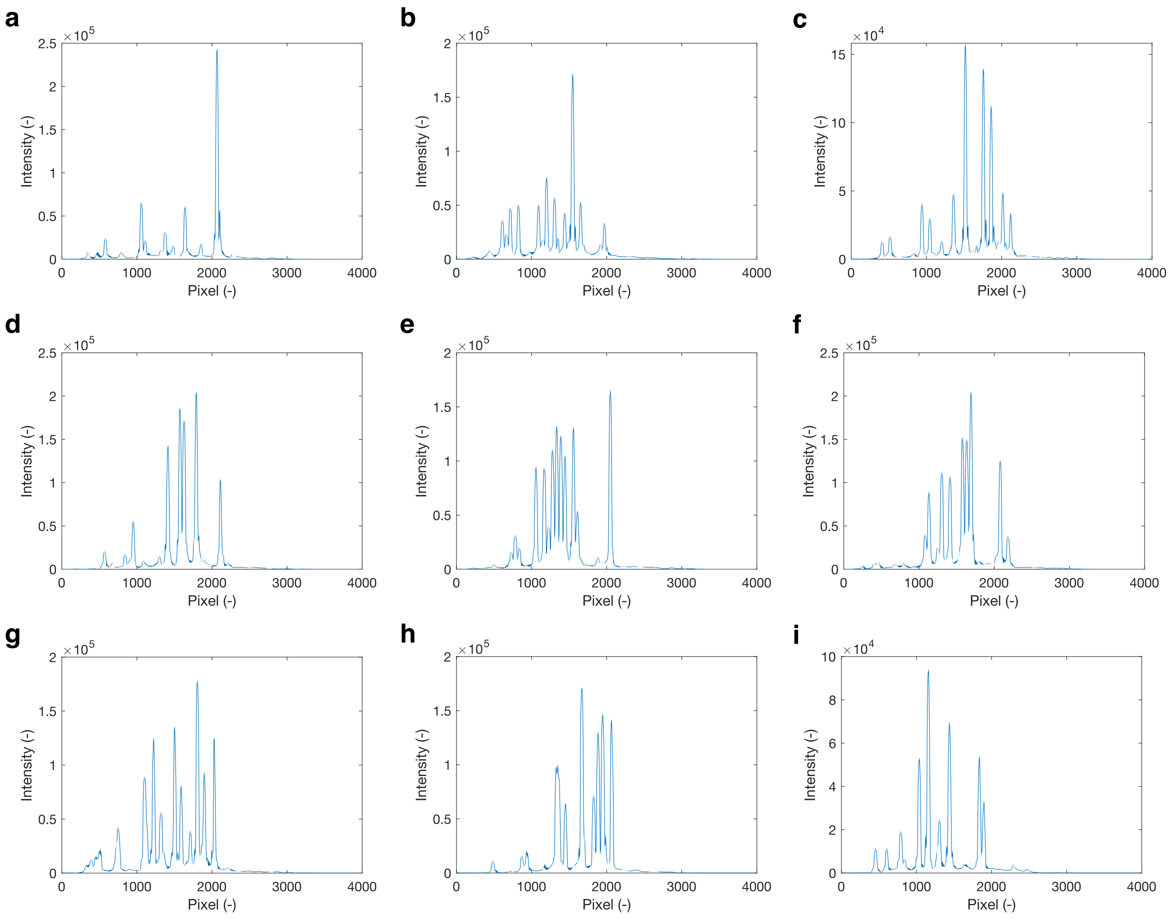


**Figure S9. Signal-to-noise ratio with HyCAP.** Summation of pixel values in each row of the cropped, grayscale images are plotted. Three representative plots selected at random are shown for Linker 1 (**a – c**), Linker 2 (**d – f**) and Linker 3 (**g – i**). According to the image processing algorithm, each peak represents a first-order dispersion band, and the area under the peak (standard width applied to be 20 pixel-rows) is total fluorescence intensity. Intensity levels between peaks is indicative of background signal which is consistent at around $2\times10^3$ au.

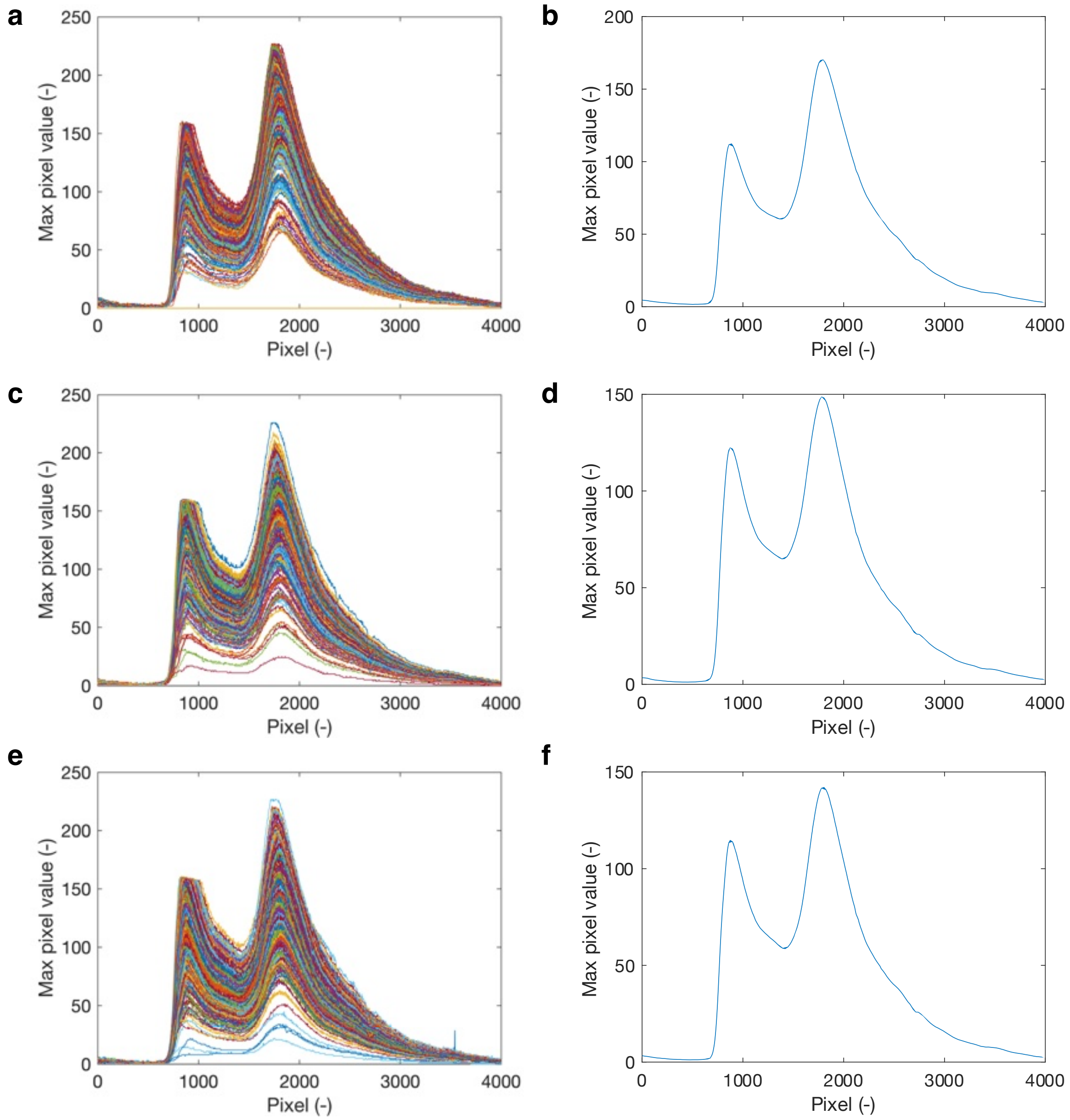


**Figure S10. Distribution of pixel values in images.** Raw (**a, c, e**) and averaged (**b, d, f**) maximum pixel values in each pixel column in grayscale images for Linker 1 (**a, b**), Linker 2 (**c, d**) and Linker 3 (**e, f**) are plotted. The exposure was adjusted such that the camera sensor was not saturated, allowing for accurate quantitation of fluorescence, and high SNR.

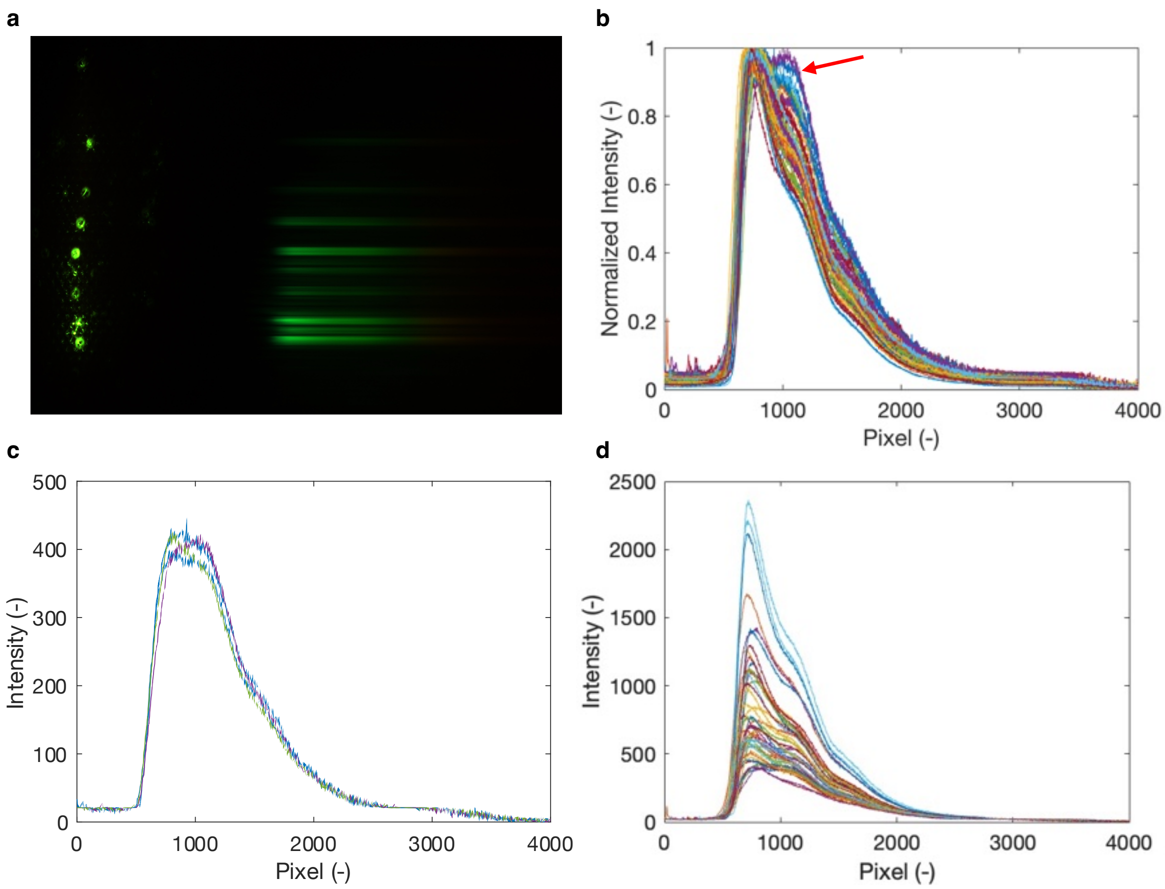


**Figure S11. mNG spectra acquisition at a shorter incubation period. a.** A representative mNG image acquired at 35× longer exposure as compared to other images is shown. **b.** Acquired spectra (49 in number) are shown. Red arrow indicates several spectra (plotted separately in **d**) with max intensity around the 1100-pixel location. These red-shifted spectra are minimal in intensity compared to all other spectra (**c**).

a

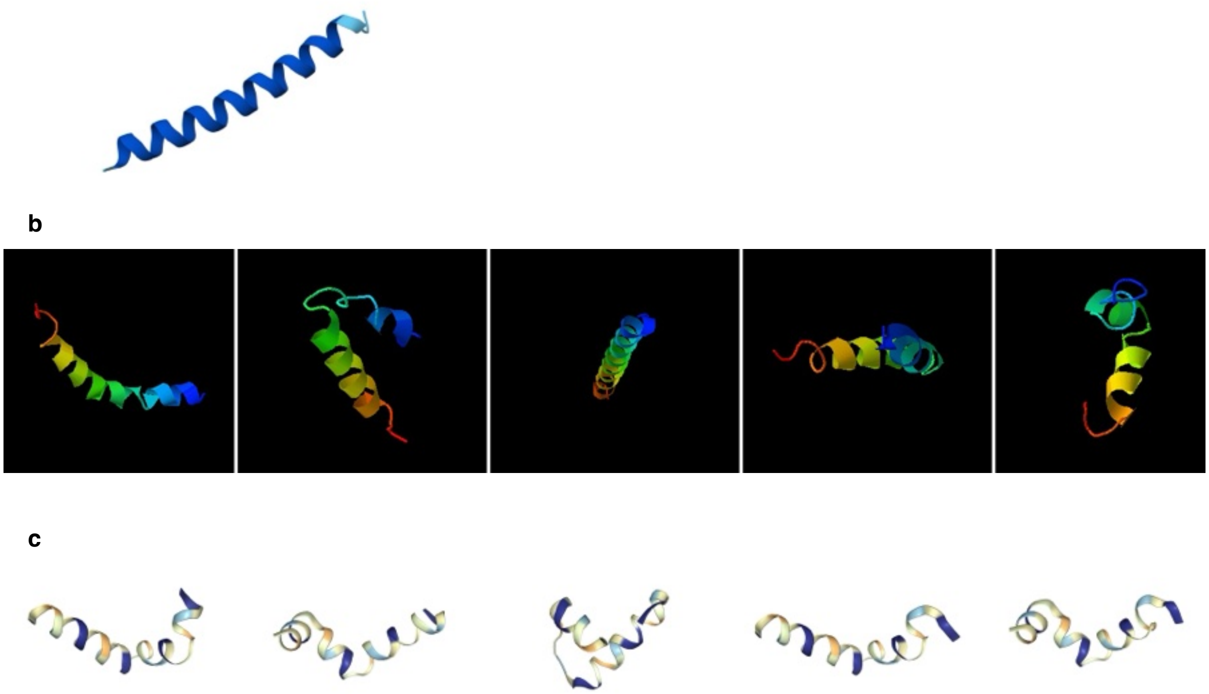

**Figure S12. Computational modelling of Linker 3.** Predicted conformations of Linker 3 as obtained from AlphaFold 3 (**a**), I-TASSER (**b**), and PEP-FOLD4 (**c**) are shown.

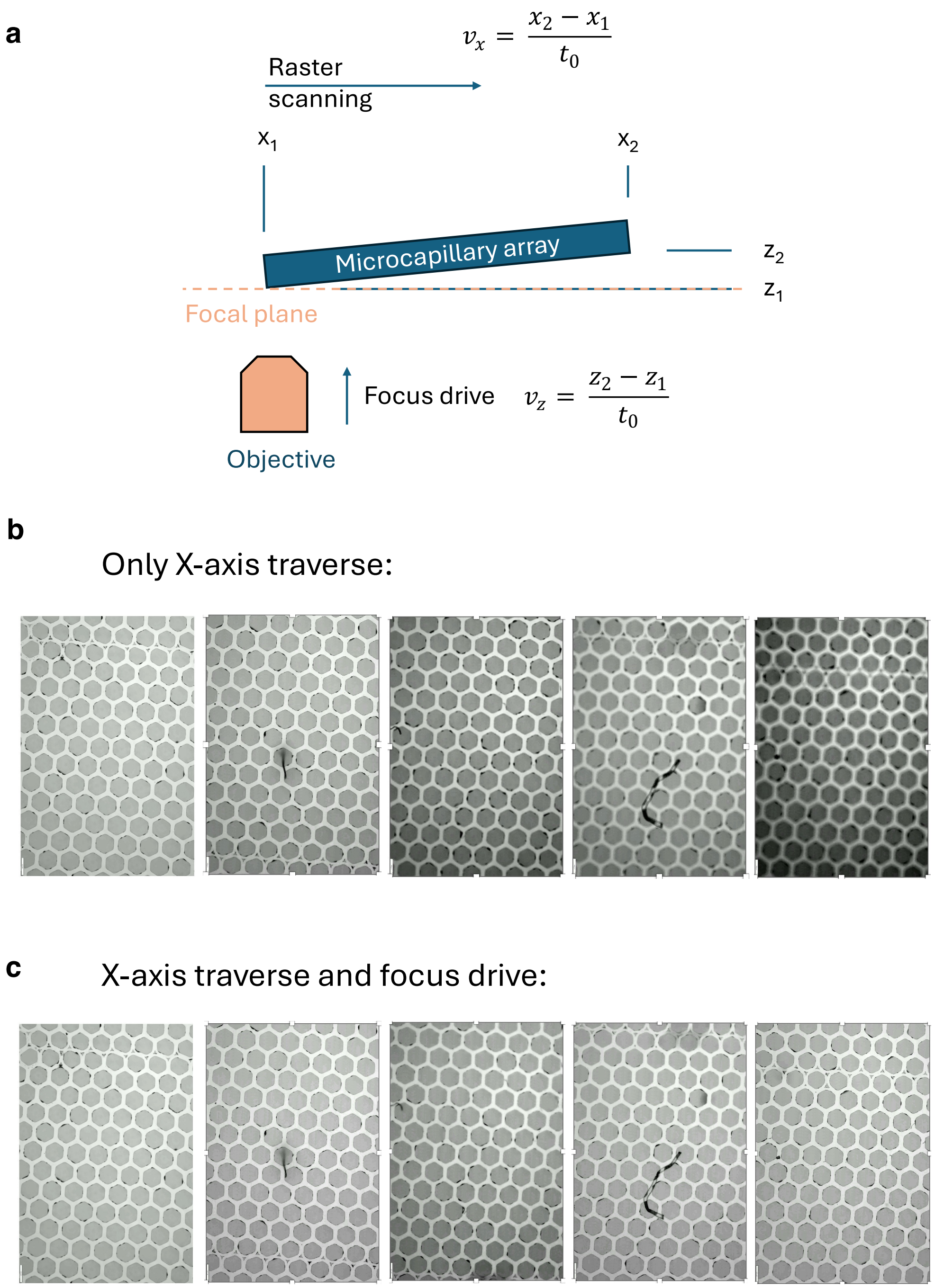


**Figure S13. Auto-focus methodology. a.** The start and end coordinates in the X and Z-axis are known. X-traverse speed is predetermined, and the corresponding traverse time-period is assigned for Z-traverse. This keeps the sample plane coincident with the focal plane of the objective. Consecutive snapshots show loss of focus with no z-traverse (**b**) vs. auto-focus (**c**).

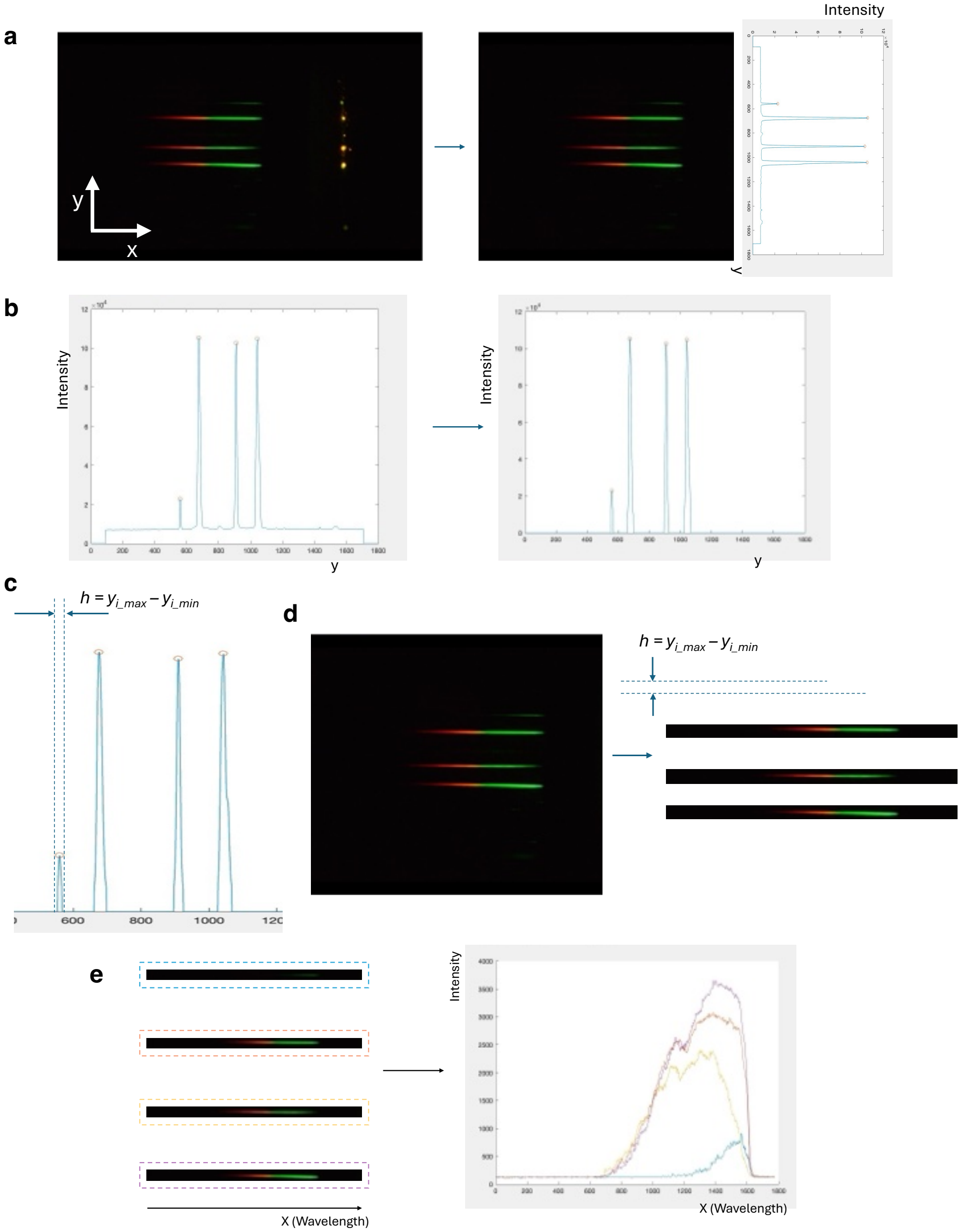


**Figure S14. Image processing algorithm, related to Methods. a.** Original image is cropped to remove zeroth-order bands, and pixel values in each row are summed and plotted to reveal intensity peaks representing positions of first-order bands. **b.** Peaks with intensity over a threshold are selected. **c.** A standard width (20 + 1 pixel rows) is assigned to all peaks. **d.** The y-coordinates of band max and min positions digitally filter each first-order bands from the ensemble. **e.** For each band, the values of all pixels in each column are summed and plotted to reveal the fluorescence emission spectrum.

| | P-value | Cohen's d |
|---|---|---|
| Linker 1-2 | 1.3105E-302 | 1.272 |
| Linker 1-3 | 3.4422E-138 | 0.81 |
| Linker 2-3 | 3.2672E-25 | 0.331 |

**Table S1. Significance and effect tests.** P-values obtained from Student's t-test, and effect sizes from Cohen's d-test are tabulated for each linker-linker combination.

##### HyCAP calibration

From theory,

$$m\lambda = dsin(\theta)$$

$$\lambda \cong d * \theta \ (small\ angle\ approx.)$$

where, $\lambda$ denotes light wavelength, $d$ is grating spacing, and $\theta$ is the diffraction angle of light.

$$d = 0.0033\ mm$$

$$\frac{1}{d} = 300\ \text{mm}^{-1}$$

Differentiating the above relation,

$$\frac{d\theta}{d\lambda} = \frac{1}{d}$$

Linear dispersion equation:

$$\text{y} = f\ tan(\theta)$$

$$y \cong f *\ \theta\ (small\ angle\ approx.)$$

where, $f$ is the focal length of the lens. Differentiating this relation,

$$\frac{dy}{d\lambda} = f * \frac{d\theta}{d\lambda} = f * \frac{1}{d} = 180\ \text{mm} * \frac{300}{\text{mm}} = 5.4\text{e}10^4$$

$$\frac{d\lambda}{dy} = 18.5\ nm/mm$$

From measurements,

3.89 um/pixel (Nikon D5300) = 0.00389 mm/pixel

14.81 pixels/nm (from slope of experimental data) = 0.00389 mm/pixel * 14.81 pixels/nm = 0.0576 mm/nm = 17.36 nm/mm.
